# A Bioinformatics Approach To Reveal Common Genes And Molecular Pathways Shared By Cutaneous Melanoma and Uveal Melanoma

**DOI:** 10.1101/2022.05.30.493940

**Authors:** Dr. Perumal Jayaraj, Seema Sen, Khushneet Kaur, Kritika Gupta, Shreya Taluja

## Abstract

**Background:** Melanomas are highly aggressive in nature and are known for metastasis and death. Melanocytes that give rise to melanomas are neural crest progenitor-cells. Our research is primarily concerned with Uveal Melanoma (UM) and Cutaneous Melanoma (CM). Although they both share same melanocytic origin, but the biology of their respective is different.

**Aim:** The aim of our study is to recognize the common differentially expressed genes (DEGs) between UM and CM.

**Methodology:** The gene expression profile was downloaded from the GEO and analyzed by GEO2R for identification of DEGs. By applying DAVID, GO and KEGG, pathway enrichment analysis was performed. PPI of these DEGs was analyzed using STRING and visualized through Cytoscape and MCODE. Further, we utilized HPA and GEPIA to obtain Kaplan-Meier Graph for survival analysis in order to assess the prognostic value of hub genes.

**Results:** We examined the UM and CM datasets and discovered three common upregulated and eight common downregulated DEGs for both the melanomas based on computational analysis. HMGCS1 and ELOVL5 were shown to be enriched in a variety of altered molecular pathways and pathways in cancer. Overexpression of HMGCS1 and ELOVL5 were linked to poor prognosis in CM.

**Conclusion:** On computational evaluation, we found that HMGCS1 and ELOVL5 were upregulated in both of these melanomas. Enrichment analysis showed that these genes are involved in cancer metabolism pathway and associated with poor prognosis in CM. However, the molecular study of these genes in UM is limited. Therefore, a better understanding of the cancer metabolism pathways should be carried to pave the way for clinical benefits.

## Introduction

Melanoma is an aggressive malignant tumor that emerges from uncontrolled proliferation of melanocytes that are pigment producing cells ^(1)^. Melanocytes are generated from neural crest progenitor cells during embryonic development giving rise to both cutaneous and choroidal cells but there are significant differences between them ^(2)^. Cutaneous melanoma (CM) originates from melanocytes that are present in the deep layer of the epidermis between them layer of basal cell, whereas in uveal melanoma (UM) melanocytes are found in the conjunctiva including all regions of the uvea (the iris, ciliary body, and choroid). Pathways engaged with melanocyte proliferation and development are known to be involved in uveal and cutaneous melanoma. Over the past decades, many studies have revealed that age, sex, genetic and phenotypic predisposition, the work environment, and dermatological diseases are among the various risk factors that lead to the development of both cutaneous and ocular melanoma ^(3)^. Among these, ultraviolet radiation has been implicated as the major risk factor for the development of cutaneous melanoma ^(4)^. It has been proposed that melanomas are caused by strong and intermittent exposure to sunlight, which does not provide time for melanocytes to synthesize melanin to protect themselves from ultraviolet irradiation (UV), resulting in DNA alterations ^(5)^. Although there is evidence to support this notion in cutaneous melanoma, the data in uveal melanoma is limited and conflicting. Although uveal melanoma (UM) and cutaneous melanoma (CM) shares common origin, but the biology of their metastasis vary differently. UM spreads through the hematogenic path and the most well-known site of metastasis of ocular melanoma in over 90% of cases is the liver. On the other hand, cutaneous melanoma can spread through both the hematogenic path and the lymphatic system, spreading to different organs such as the lung, lymph nodes, brain, soft tissue as well as different parts of the skin ^(6)^. Therefore, cutaneous melanoma is known for its remarkable ability to metastasize and is considered more aggressive than uveal melanoma of similar histogenesis.

The etiology of both uveal and cutaneous melanoma is complex and heterogeneous. In CM, B. Uzdensky et al., 2013 stated that mutations were observed in the Ras/Raf/MEK pathway, which is a key regulator of cell proliferation and survival ^(7)^. Signature mutations in Ras or Raf proteins promoted cell proliferation, tumor invasion, and metastasis indefinitely. Permanent cell proliferation, apoptosis resistance, and malignant transformation are all caused by mutations in the NRAS and BRAF genes. The activating BRAF mutations are found to be important in the pathophysiology of melanoma. It has been demonstrated that mutated B-Raf irreversibly activates ERK and increases proliferation ^(7)^. Additionally, various studies have also revealed that cutaneous melanoma patients frequently harbor alterations in NF1, TP53, and CDKN2A, where CDKN2A mutations were susceptible in both CM and UM ^(8)^. Katopodis et al, 2021 specified one of the key molecular pathways observed in uveal melanoma is PI3K/Akt/PTEN which is involved in cell proliferation ^(9)^. BAP1, one of the most significant mutations is recognized in uveal melanoma ^(10)^, as well as mutations in GNAQ and GNA11, are the most frequent mutations observed in UM. At the cellular level, cutaneous and uveal melanomas share certain molecular profile as several studies have indicated that in both melanoma types, the mitogen-activated protein kinase (MAPK) pathway is involved and it regulates melanogenesis. In addition, the phosphatidylinositol 3-kinase (PI3K)/protein kinase B (Akt) pathways that regulate cell growth, differentiation, migration, and survival, as well as angiogenesis and metabolism are also known to undergo dysregulation leading to genetic alterations and poor treatment outcomes in both CM and UM. Also the microphthalmia (MITF) transcription factor which is critical to the growth development and progression of melanocytes and melanoma ^(11)^ is known to be biochemically targeted by the c-kit signalling pathway implying a link to proliferation ^(12)^. Ortega et al, 2020 have revealed that mutations in the telomerase reverse transcriptase (TERT) gene and EIF1AX have been found to be common with both cutaneous melanoma and ocular melanoma ^(3)^.

There has been a revolution in understanding the molecular complexity of melanoma biology and there have been tremendous advancements in treatment over the previous decade for CM but the same for UM has been gradual. In this present study, the utilization of computational networking analysis along with various online bioinformatic tools has given us means to investigate further about the relationship between uveal melanoma and cutaneous melanoma. The aim was to recognize the common differentially expressed genes (DEGs), dysregulated genes, the cancer pathways, and the metabolic pathways which may be common to both these melanoma tumor types.

## Material and Methods

### Data procurement

The gene expression profile datasets used in this investigation were collected from NCBI’s database-Gene Expression Omnibus (GEO) [https://www.ncbi.nlm.nih.gov/geo/]. The database yielded a total of 43,707 series related to human cutaneous melanoma and 1227 series related to human uveal melanoma. The gene expression profiles GSE15605 ^(13)^ and GSE44295 ^(14)^ were selected after careful contemplation for cutaneous and uveal melanoma respectively. Amongst the two, GSE15605 was based on platform GPL570 [HG-U133_Plus_2] Affymetrix Human Genome U133 Plus 2.0 Array while GSE44295 was based on the platform GPL6883 Illumina HumanRef-8 v3.0 expression beadchip. All of the data is publicly available online, and no trials on humans or animals were done by any of the authors for this study. As public records were evaluated, no ethical approval was required for this study (Figure 1).

**FIGURE 1:**
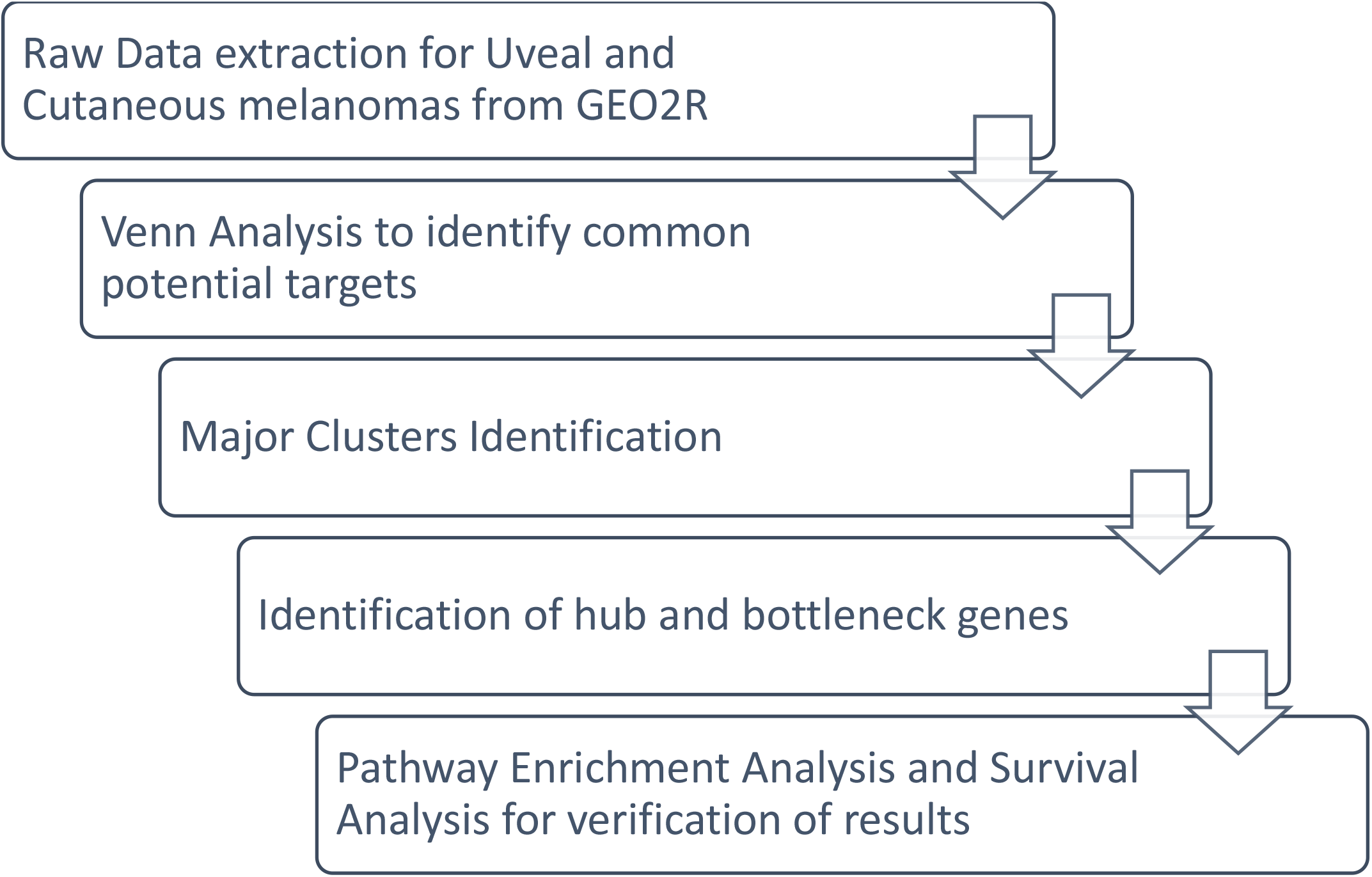
Workflow of the study.

### Data Processing of DEGs

An online freely available tool, GEO2R (www.ncbi.nlm.nih.gov/geo/geo2r/), was used to identify the DEGs for uveal and cutaneous melanoma by comparing the tumor with normal specimens. DEGs were defined as genes that satisfied the cut-off criterion of adjusted P0.05 and | logFC|>1.0. For each dataset, statistical analysis was done, and the Venn diagram online tool (bioinformatics.psb.ugent.be/webtools/Venn) was employed to find the common DEGs shared by both uveal and cutaneous melanomas.

### PPI Network Construction and Module Evaluation

Establishment of PPI Networks and Module Analysis. Using the STRING web-based tool (v11.0, 3) (https://string-db.org/) to analyse the association among the DEGs from the obtained datasets, we built a PPI (protein-protein interaction) network. The network was visualised using Cytoscape software ^(15)^ and major gene clusters were identified. Cluster two, with maximum number of genes was selected for further analysis, Centiscape plugin on Cytoscape was used to identify the Hub genes. Genes with maximum MCC score were regarded as the Hub genes and further studied.

### Gene ontology analysis

Gene Ontology is a frequently used approach for linking genes, gene products, and sequences to their biological processes. Gene functions are divided into three categories: biological process (BP), molecular function (MF), and cellular component (CC). KEGG (Kyoto Encyclopedia of Genes and Genomes) is a database which can be used to understand genomic sequences and other high-throughput data in a biological context. Both these analyses were done using the ShinyGO database ^(16)^ which is a bioinformatics data repository with genomes and pathways information for over 315 plant and animal species, based on Ensembl, Ensembl plants, Ensembl Metazoa, STRING, and KEGG databases. P value <0.05 was considered significant.

### Survival analysis of key genes

The Human Protein Atlas ^(17)^ is an open access software with information in protein upon tissue, brain section, single cell type, tissue cell type, metabolism etc. We used HPA database to obtain the Kaplan Meier plots of the key genes, wherein p values <0.05 were considered significant. The Km plots were used to analyze their role as prognostic markers for cancer.

### Immunohistochemical evaluation

The Human Protein Atlas (HPA, https://www.proteinatlas.org/) database was used to obtain immunohistochemical staining to verify the protein expression level of key genes in cutaneous melanoma. Although data for uveal melanoma was not available in any of the free databases.

## Results

### Identification of common genes

GSE15605 and GSE44295 (Table 1) are two datasets retrieved from the GEO repository that were then subjected to data analysis in accordance with our inclusion criteria. The P <0.05 and |logFC|>1 criterion was used to retrieve DEGs from the GEO database. The overlap between the two datasets is seen in the Venn diagram (figure 2). Four hundred and ninety-two DEGs were significantly similar in expression between the two groups and were included in the study.

**Table 1:**
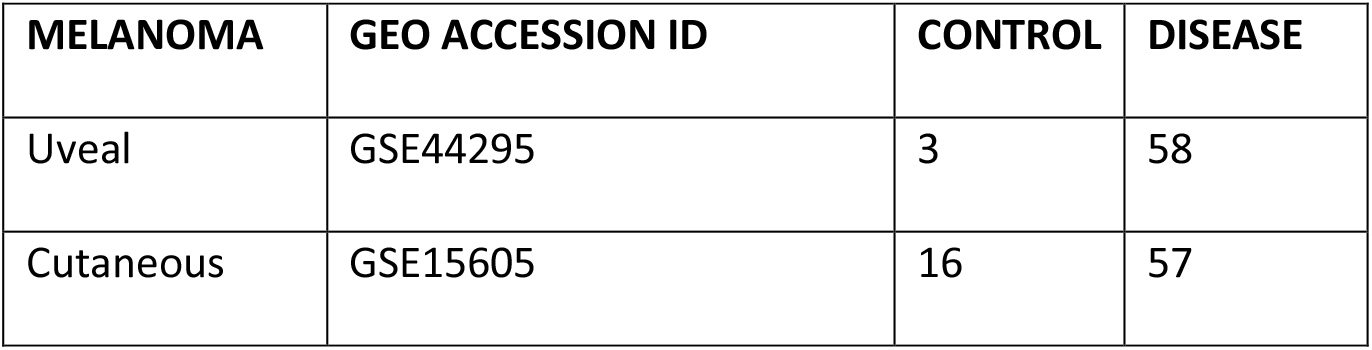
GEO datasets chosen for the study obtained from NCBI GEO2R database.

**FIGURE 2:**
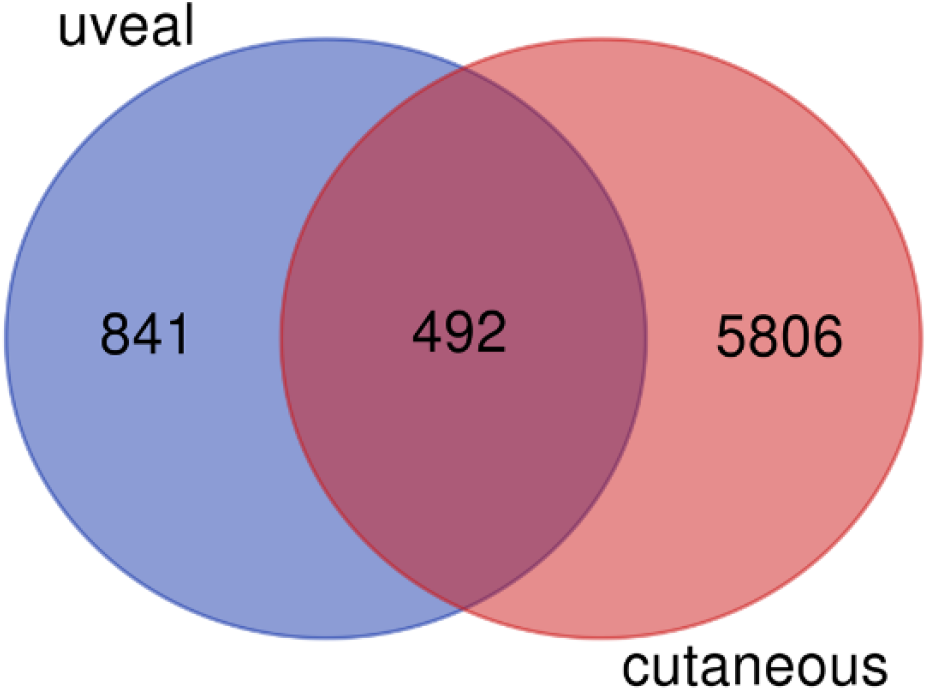
Venn diagram showcasing 492 common differentially expressed genes between uveal and cutaneous melanomas.

### PPI Network Construction and Module Evaluation

Employing STRING online tool, the DEG PPI network was constructed (Figure 3). Cluster 2, was identified using MCODE in Cytoscape (Figure 4A) and it displayed significant level of common DEGs (Table 2). It was considered as the most important module and was selected for further analysis. Further 3 upregulated and 8 downregulated genes (Figure 4B) were identified based on their log Fc values. Following calculations by centiscape, screening of HUB genes using maximum MCC score was done. (Table 3)

**FIGURE 3:**
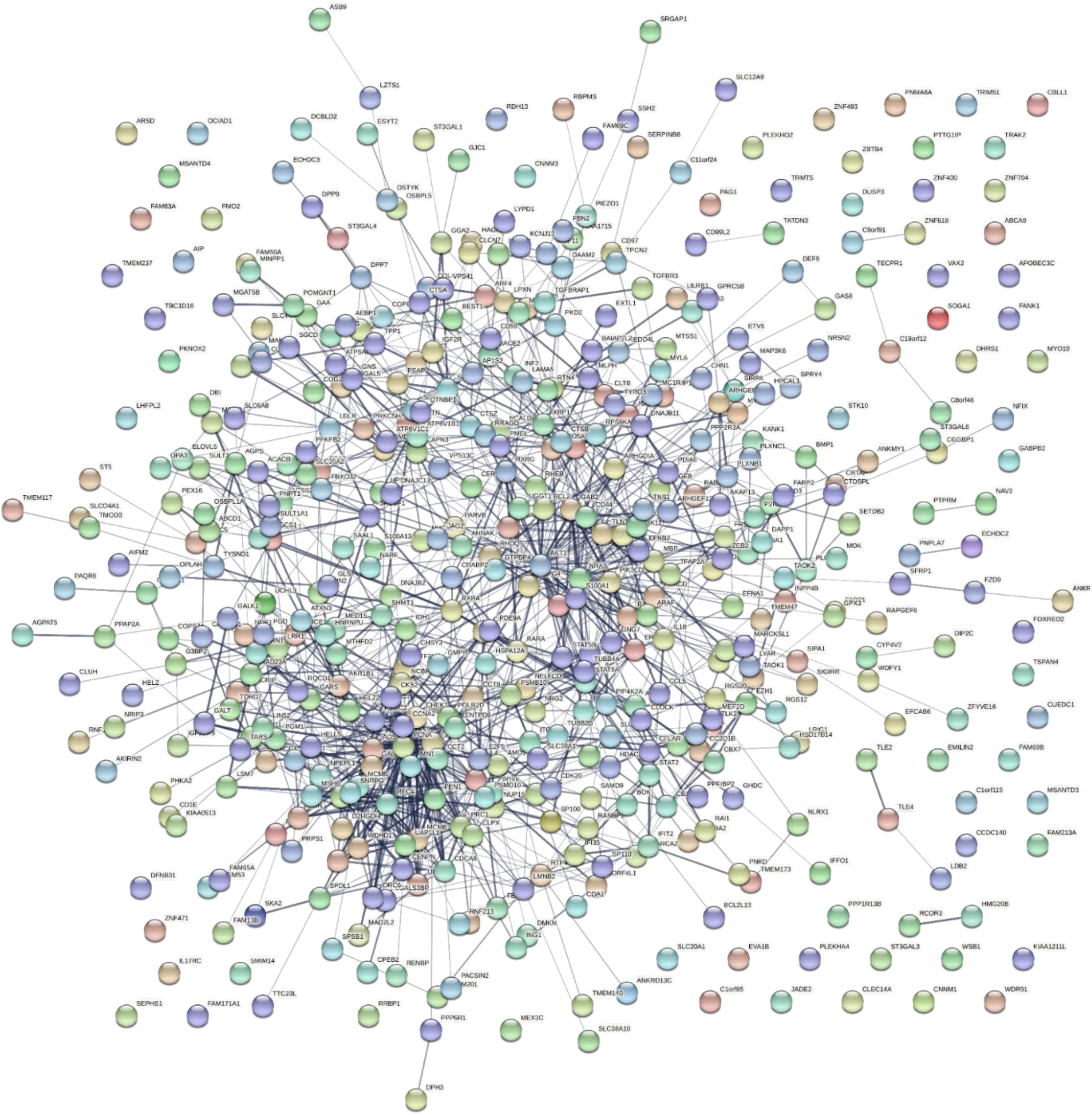
Protein-protein Interactions for the 492 targets as obtained from the STRING Network.

**FIGURE 4:**
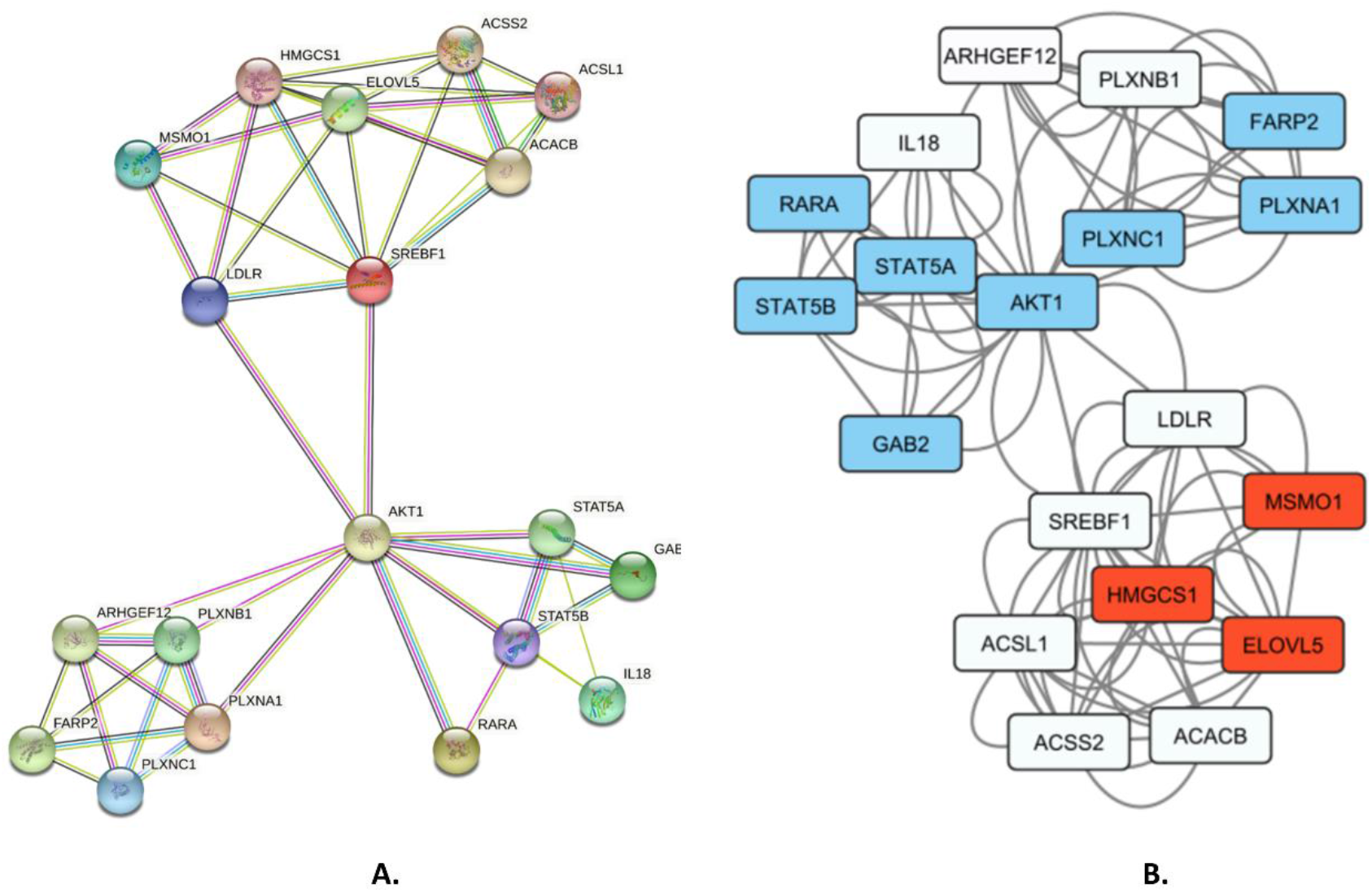
A. STRING PPI Network as obtained for the genes present in the Cluster-2. **B. Cluster 2 genes, where blue color denotes downregulated genes and red color denotes the upregulated genes in both uveal and cutaneous melanomas**

**Table 2:** Upregulation and downregulation of cluster 2 genes in uveal and cutaneous melanomas.

| GENE | UVEAL | CUTANEOUS |
| --- | --- | --- |
| CLUSTER 2 |  |  |
| RARA | down | down |
| STAT5B | down | down |
| ARHGEF12 | down | up |
| STAT5A | down | down |
| AKT1 | down | down |
| PLXNB1 | down | up |
| HMGCS1 | up | up |
| MSMO1 | up | up |
| ACSS2 | down | up |
| LDLR | up | down |
| IL18 | down | up |
| FARP2 | down | down |
| ACACB | down | up |
| ELOVL5 | up | up |
| PLXNC1 | down | down |
| GAB2 | down | down |
| ACSL1 | down | up |
| SREBF1 | down | up |
| PLXNA1 | down | down |

**Table 3:**
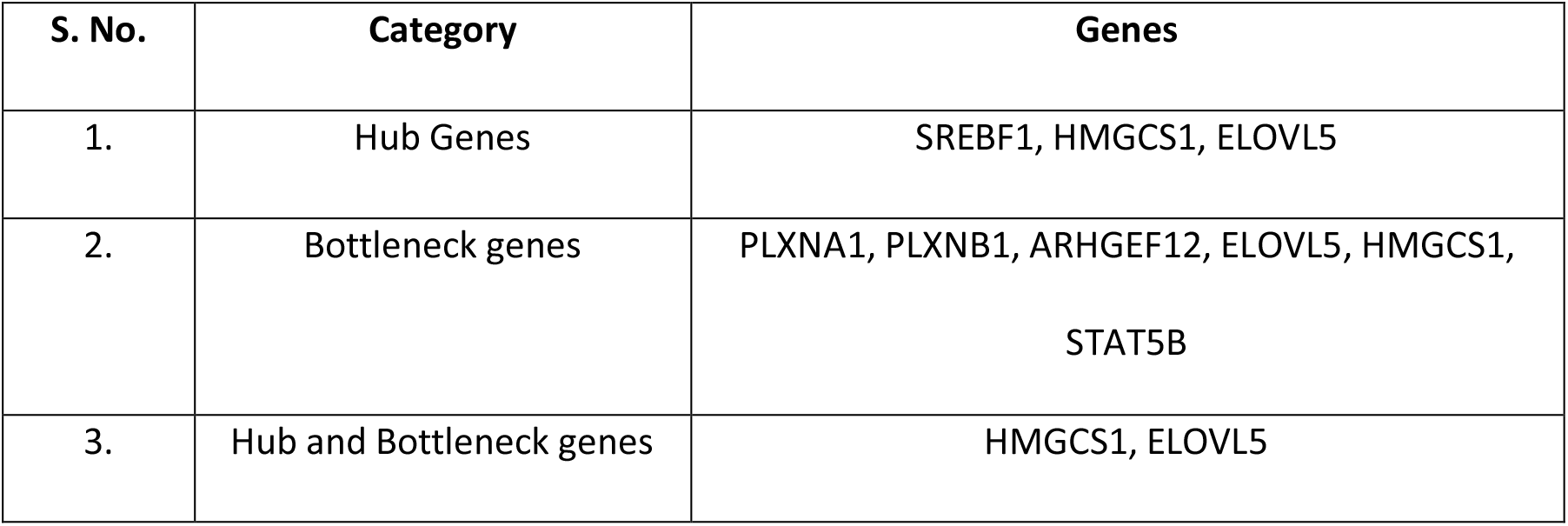
Hub and Bottleneck genes identified using degree and betweenness filter respectively.

### Functional and Pathway Enrichment

Database for Annotation, Visualization and Integrated Discovery (DAVID) was used to perform functional and pathway enrichment analysis on Cluster 2 DEGs in order to better understand their biological categorization. According to the results of the GO analysis, DEGs were primarily involved in metabolic pathways such as fatty acid metabolism, lipid biosynthetic and metabolic process, semaphoring-plexin signalling pathway (Figure 5A)

**FIGURE 5:**
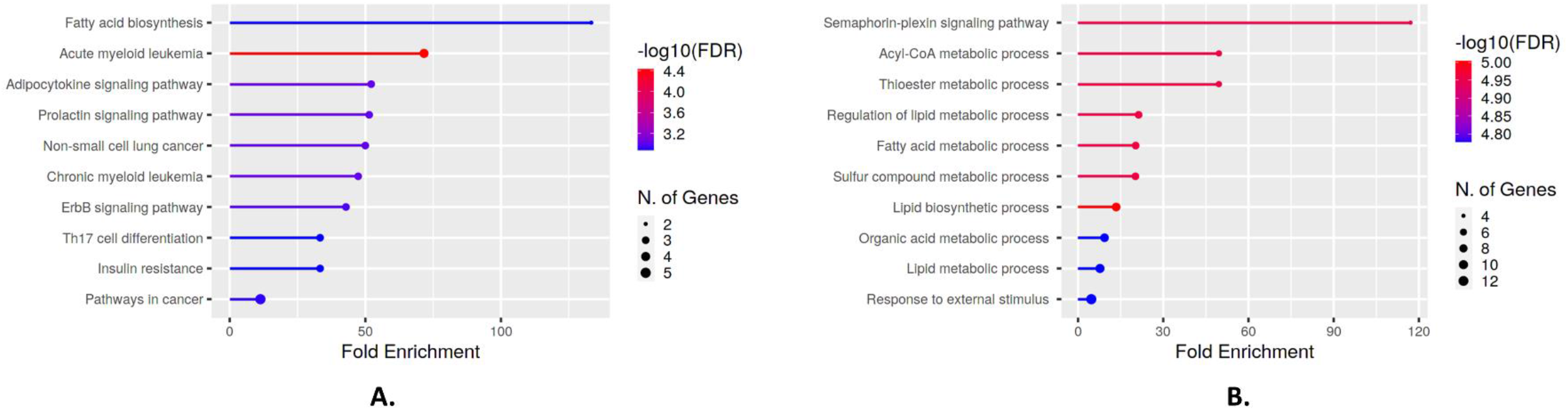
A. Gene Ontology for the cluster 2 genes as obtained using ShinyGO (FDR=0.01) **B. Top 10 KEGG Pathways obtained for the cluster-2 genes**

Furthermore, the findings of the KEGG pathway study revealed that DEGs were shown to be prominent in the cancer metabolism pathway, fatty acid biosynthesis, and ErbB signalling pathway, Th 17 cell differentiation among other molecular pathways (Figure 5B).

### Survival Analysis

The Kaplan-Meier Plot was used for survival analysis to evaluate the prognostics significance of hub genes in cutaneous melanoma. High expression of HMGCS1 and ELOVL5 was shown to be related with a poor prognosis in cutaneous melanoma. (Figure 6A,6B).

**FIGURE 6:**
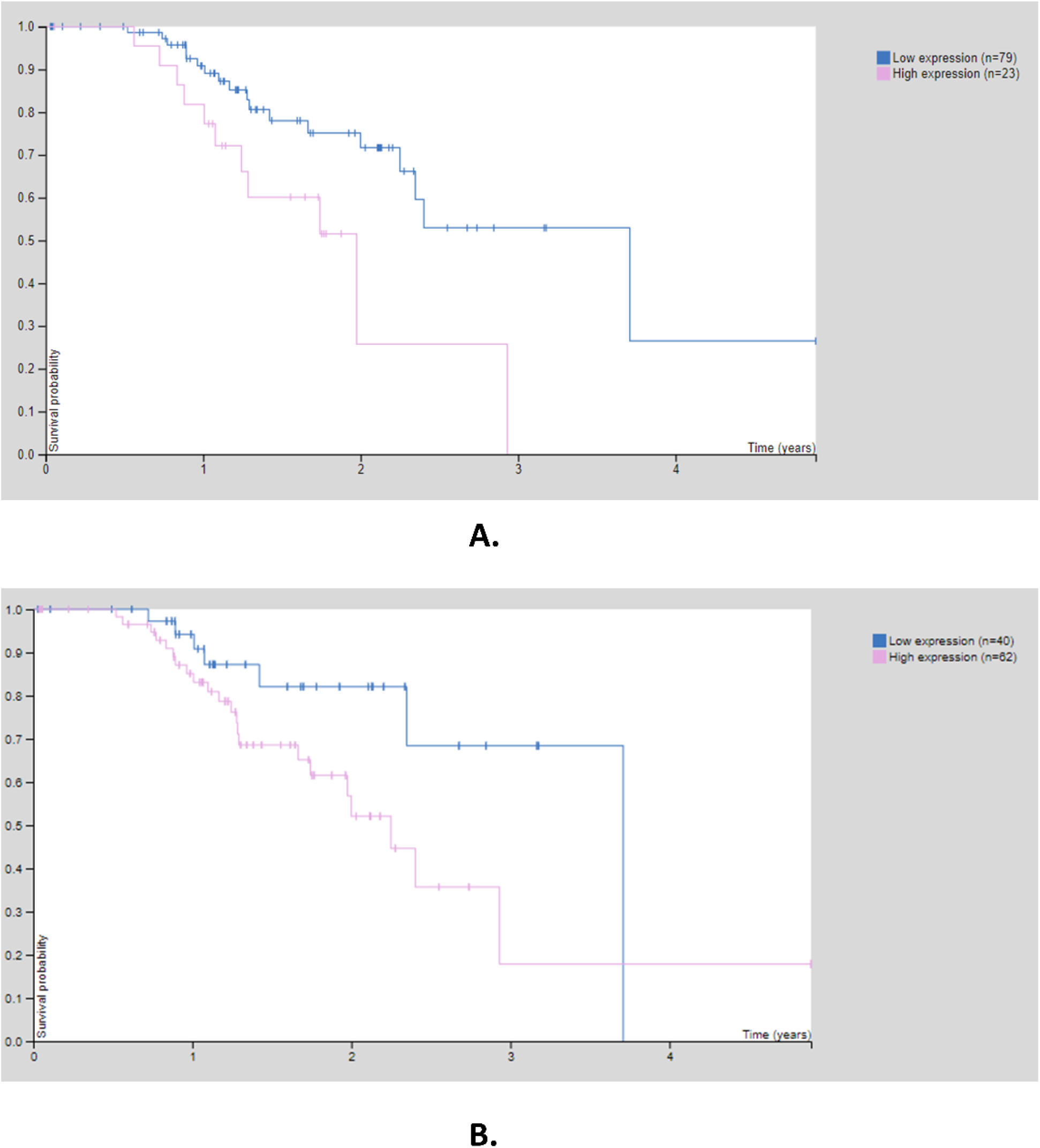
Km Plots for A. HMGCS1 and B. ELOVL-5 expression in melanoma obtained from the Human Protein Atlas showcasing regressed survival rates in patients with high expression levels of these genes.

**FIGURE 7:**
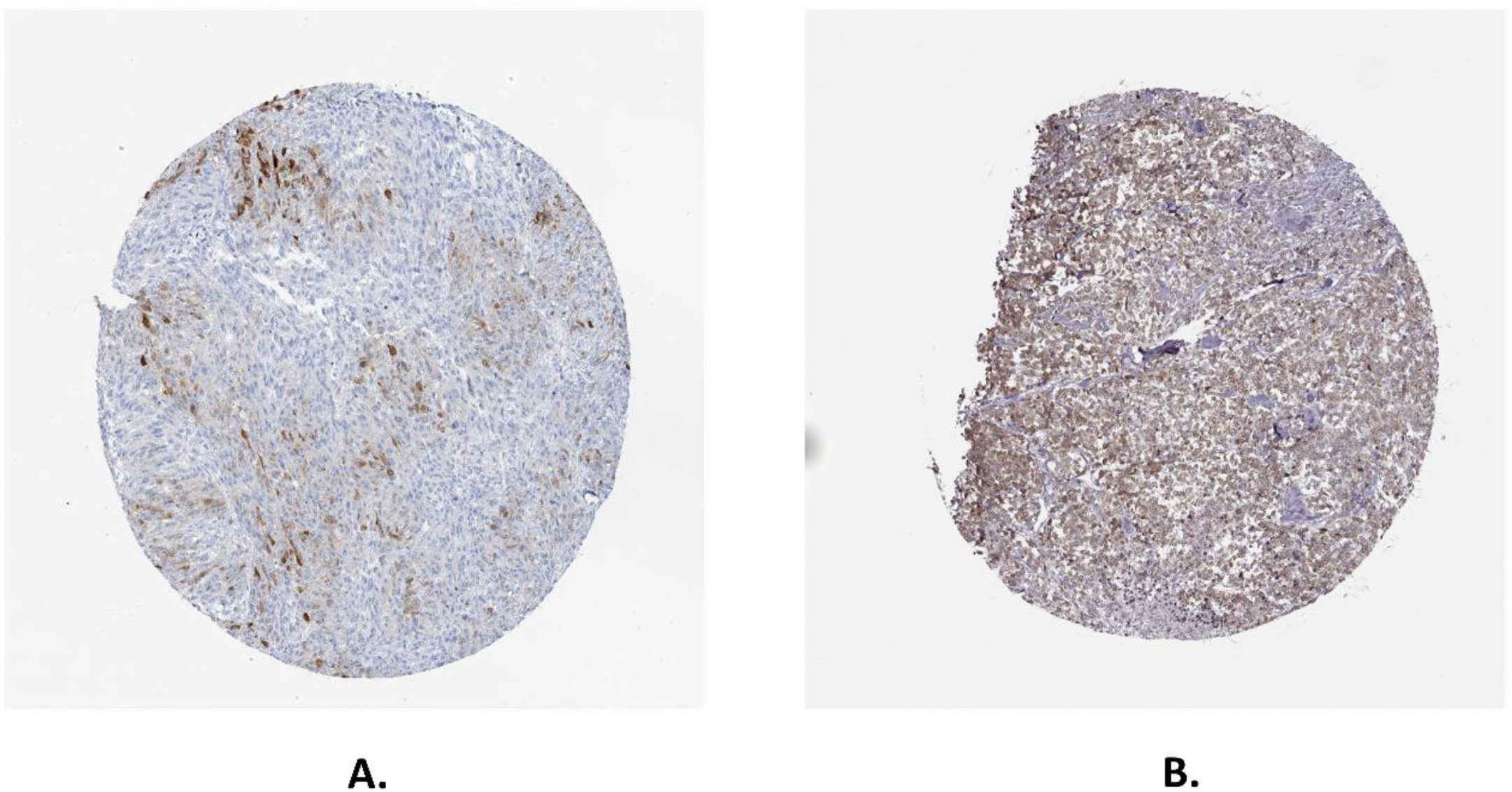
(A) Strong cytoplasmic and membranous Immunohistochemical expression of HMGCS1 in case of Cutaneous melanoma. (B) Cytoplasmic over expression of ELOVL5 in a representative case of cutaneous melanoma.

### Immunohistochemical expression

ELOVL5 AND HMGCS1 immunohistochemical staining was shown to be positive in cutaneous melanoma tissues. For HMGCS1, Strong cytoplasmic and membranous expression was observed. However, Cytoplasmic over expression of ELOVL5 was noted.

## Discussion

Melanoma is a malignancy of melanocytes and is known for its remarkable ability to metastasize rapidly. Cutaneous Melanoma (CM) and Uveal Melanoma (UM) are two common forms of melanoma that arise from the same embryonic origin but the etiopathogenesis and biological characteristics of these melanomas, however, are highly distinct. Despite the fact that they both have same melanocytic origins, the biology of their respective metastasis is different. The goal of our research was to find common differentially expressed genes (DEGs) in UM and CM.

Using Bioinformatics approach, we identified 492 common DEGs that formed a close network in STRING. The reconstructed network in Cytoscape was used to filter significant sub-network modules using the MCODE plugin, and module two was chosen because it had the highest number of shared upregulated and downregulated genes. On performing DAVID and KEGG databases, an analysis of pathway enrichment found that 106 metabolic processes and pathways were implicated in cluster 2. Furthermore, it was notably rich in metabolic processes such as acyl-COA, lipid regulation, fatty acid metabolism, and lipid biosynthetic process as well as Pathways like erbB signalling Pathway, Th17 cell differentiation, and cancer pathways. Subsequently, from cluster four, Hub genes namely HMGCS1, ELOVL5, SREBF1, and AKT1 were filtered based on the greatest MCC score using the Centiscape plugin, indicating a path to examine the complex interaction of these genes. Further, out of these, HMGCS1 AND ELOVL5 were found to be significantly upregulated. Several studies have already indicated HMGCS1 and ELOVL5 to be oncogenic in nature and to be involved in various cancer and metabolic pathways.

3-hydroxy 3-methylglutaryl-CoA synthase 1, (HMGCS1) is known to be potential regulatory node of the mevalonate pathway which is further involved in tumorigenesis by positively regulating the cell proliferation, migration, invasion, and metastasis of tumor cells ^(18)^. HMGCS1 overexpression has been observed in various cancers like breast cancer, gastric cancer, renal cancer, colon cancer, and prostate cancer. Shengzhou et al, 2020 has demonstrated that the suppression of HMGCS1 could completely reduce EGF-induced proliferation of colon cancer cells indicating HMGCS1 as a novel biomarker ^(19)^. Elongation of very long chain fatty acid-like (ELOVL5) is necessary for the production of C20-22 PUFAs. It plays significant function in the synthesis of fatty acid elongation ^(20)^ influenced cell migration, cell– cell interactions, and MMP’s synthesis ^(21)^.Substantial evidences suggest that ELOVL5 expression is commonly deregulated in cancer. It has been found to be overexpressed in hepatocellular carcinoma^(22)^ lung cancer ^(23)^, and breast cancer^(24)^. Nevertheless, many experimental models have demonstrated ELOVL5 to be extremely low in certain cancers ^(25)^, therefore, further research on the understanding of this gene in cancer metastasis is warranted.

Therefore, it can be suggested that the upregulation of ELOVL5 and HMGCS1 in several carcinoma might play an important role in the pathogenesis of C, thereby, indicating a key role for these novel genes as potential therapeutic targets.

Our study reveals a network-based strategy to screen genes and understand multiple metabolic pathways responsible for both CM and UM using data mining and bioinformatic analysis approaches. However, due to limited number of patients and difficulty in access to tissues, progress in researching and identifying genes for uveal melanoma has been limited. The study of cell differentiation pathways and genes, as well as numerous biochemical processes, can assist in the early diagnosis andtreatment of melanoma. Therefore, hub genes implicated in the advancement of CM and UM through our study may be investigated further for aid in the development of prognostic and therapeutic techniques to cure UM using concepts similar to those used in CM.

